# Foraging egg parasitoids benefit from the presence of potential competitors

**DOI:** 10.64898/2025.12.22.695894

**Authors:** Prabitha Mohan, Vanshika Pal, Abhyudai Singh, Saskya van Nouhuys

**Author notes:** Email address: Prabitha Mohan^1^, Vanshika Pal^1,3^, Abhyudai Singh^4^.

## Abstract

Competition is generally considered a negative interaction that lowers the fitness of individuals. Studies of competition and fitness traits in parasitoids have primarily focused on interference competition, with less attention paid to parasitoids under exploitative competition. Theoretical studies predict that interference competition will decrease the resource utilisation of parasitoids, while exploitative competition will not. In our study, we investigated the impact of intraspecific exploitative competition among parasitoids on fitness traits, including resource utilisation, the percentage of offspring that emerged from parasitised eggs, and the percentage of female offspring produced from parasitised eggs. We investigated whether these fitness traits were influenced by the presence of conspecific foraging females, as well as by parasitoid and resource densities. Since we were studying the effect of exploitative competition, we used *Trichogramma chilonis*, a non-aggressive egg parasitoid. *Corcyra cephalonica*, a stored grain pest, was used as the host. We found that *T. chilonis* resource utilisation increased in the presence of conspecifics. However, the density of those conspecifics and the resource density did not affect resource utilisation. The presence of conspecific foraging females did not affect the rate of offspring survival. However, at a lower resource density, offspring survival was higher than under resource-abundant conditions. We explored this counterintuitive pattern using a simple mathematical model, which shows that superparasitism, occurring at high parasitoid density, can lead to high emergence. This is because, given only some progeny survive, the chance of parasitoid emergence increases with the number of eggs laid in the host. The secondary sex ratio of the progeny was unaffected by the presence of conspecifics or by parasitoid or resource densities. We conclude that for *T. chilonis*, the presence of competitors has a positive effect. Its high performance under competitive conditions may contribute to its efficiency as a biological control agent.

## Introduction

Competition is an important biotic factor that is considered a key driver of population dynamics, patterns of species coexistence, and community structure (Bernhardt et al., 2020). Competition can impact the behaviour and life history traits of organisms (Kaplan & Denno, 2007; Wauters et al., 2019; Ode et al., 2022) and is generally considered a negative interaction because of its negative fitness consequences for the involved parties (Cody et al., 1975; Forsman et al., 2002; Koutsidi et al., 2024). For instance, under competition with the frog *Pelodytes punctatus*, the bullfrog *Bufo bufo* increases its foraging activity, develops slowly and its size at metamorphosis declines (Richter-Boix et al., 2004) The intensity of competition is affected by the density of competitors and the availability of resources (Johnson et al., 2004; Hasegawa & Yamamoto, 2009; Pekkonen et al., 2013; Riley & Dybdahl, 2015; Wang & Callaway, 2021; Tao et al., 2024). For instance, density-dependent competition as a result of limited food availability in mosquitoes can cause increased juvenile mortality, delayed pupation, and reduced adult longevity (Agnew et al., 2002; Than et al., 2020).

Parasitoid wasps experience intense competition compared to other higher trophic-level animals because, unlike predators, individual parasitoids rely on a single host for all the resources necessary for their development (Ode et al., 2022). Resource competition happens at various stages of a parasitoid’s lifecycle, including both extrinsic competition between adult females for oviposition and intrinsic competition between juveniles inside or on a host for nutrition (Harvey et al., 2013; Cusumano et al., 2016; Ode et al., 2022). Both extrinsic and intrinsic competition can happen between individuals of the same species (intraspecific competition) or between individuals of different species (interspecific competition).

Competition affects several fitness traits of parasitoids, including developmental time, offspring survival, and body size (Harvey et al., 2013, 2019; Visser et al., 2014; Cusumano et al., 2015). Intrinsic competition among multiple parasitoid larvae within a host can cause resource limitation, leading to long development time and small offspring size (Harvey et al., 2009). Under low resource availability, interspecific competition can cause offspring mortality through superparasitism (Xu et al., 2013). However, in some situations, superparasitism can be advantageous at the individual or population level (van Alphen & Visser, 1990). The presence of conspecific adult females can cause a shift in the offspring’s primary sex ratio (the sex of eggs laid), increasing the proportion of males. This is found where mating occurs locally (Hamilton, 1967; King, 1993; King & Seidl, 1993), or when resource limitation leads to lower resource availability for offspring development, since males are generally smaller than females (Brodeur & Boivin, 2004).

Adult female parasitoids foraging for hosts behave differently depending on the density of conspecifics and the density of host resources available (Xu et al., 2013; Couchoux & van Nouhuys, 2014; Cusumano et al., 2016; De Rijk et al., 2016). Studies of egg parasitoids show that the number of eggs parasitised by a single female often increases with increasing host and parasitoid density (Kfir, 1981, 1983). Some parasitoids reduce their host patch exploitation time, leaving the patch early in the presence of conspecifics to avoid direct competition (Wajnberg et al., 2000; Robert et al., 2016; Kishani Farahani et al., 2019). For example, the patch exploitation time of a pupal parasitoid, *Pachycrepoideus vindemmiae,* decreases with increasing competitors to avoid direct competition (Goubault et al., 2005).

Competition among adult parasitoid females can be divided into interference or exploitative (Cusumano et al., 2016; Ode et al., 2022). Under intraspecific interference competition, parasitoid adult females may either chase away their competitors or physically combat them to guard their host resources (Batchelor et al., 2005; Nakamatsu et al., 2009). Under intraspecific exploitative competition, they will alter their foraging activity spatially or temporally to reduce the direct adverse effects of competition (Ode et al., 2022). Theoretical models predict that the rate of resource use of parasitoids decreases under interference competition, and remains unaffected under exploitative competition (Robert et al., 2016). However, empirical evidence of behaviour during exploitative competition and its consequences is scarce (Robert et al., 2016). In this study, we investigated the impact of intraspecific exploitative competition on resource utilisation and two other performance traits of parasitoids. We subjected the wasps to two drivers of competition: resource availability (host egg density) and conspecific density. We hypothesised that individual wasps would parasitise more eggs in the presence of conspecifics. This strategy may help offset the potential loss of their offspring due to superparasitism (Ives, 1989). Consequently, we expect the number of eggs parasitised per female parasitoid will increase with increasing parasitoid density and decreasing resource density. We also expected the secondary sex ratio to be less female-biased under an increasing number of conspecifics and lower resource density.

The non-aggressive egg parasitoid *Trichogramma chilonis* Ishii (Trichogrammatidae, Hymenoptera) is used for the study. *Trichogramma chilonis* is a biocontrol agent widely used against various pest Lepidoptera species such as *Plutella xylostella, Trichoplusia ni,* and *Crocidolomia pavonana* (Miura et al., 1994; Miura & Kobayashi, 1998; Singhamuni et al., 2015). Experienced *T. chilonis* females can discriminate parasitised from unparasitised eggs using their antennal perception and tend to lay eggs on unparasitised host eggs (Miura et al., 1994; Wang et al., 2016). We used *Corcyra cephalonica* (Pyralidae: Lepidoptera) eggs as the host for the study. One *C. cephalonica* egg parasitised by *T. chilonis* usually produces a single offspring (Chowdhury et al., 2016; Manisha et al., 2020). The number of eggs parasitised per female was used to quantify resource utilisation per female under different competitive conditions. The percentage of offspring that emerged (offspring survival) and the percentage of female offspring out of the total offspring that emerged were used as performance traits. Resource utilisation and performance traits were observed under densities of one, two, four, and eight females to study the effect of adult parasitoid densities. To study the impact of resource density, resource utilisation and performance traits were observed under Resource Abundant (RA) and Resource Constant (RC) conditions (Fig. 1). In RA conditions, the resource density was increased with increasing parasitoid density. In contrast, in RC conditions, the resource density was kept constant irrespective of the parasitoid density. We expected that resource utilisation would be higher in the RC condition than in the RA condition due to higher competitive pressure. Finally, since we found that the rate of emergence from parasitised eggs increased with the density of females under RC conditions, we then made a simple probabilistic model to explore the idea that this could be explained by survival under superparasitism, which increases when resources are limited.

**Figure 1:**
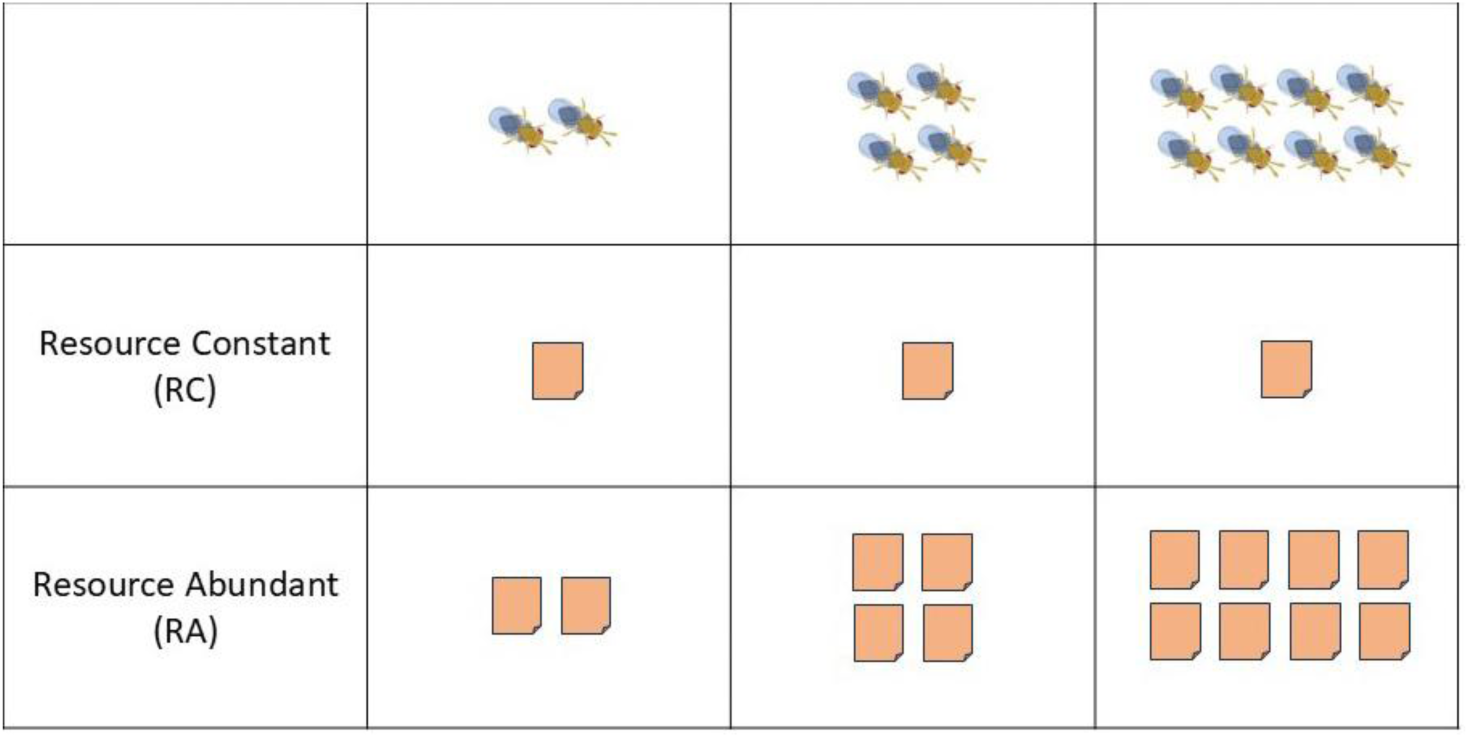
Graphical representation showing resource distribution under two different resource densities: Resource Abundant **(RA)** and Resource Constant **(RC)** conditions. Each egg card (orange square) contains 500-600 *C. cephalonica* eggs.

## Methods Insect Rearing

The initial cultures of the parasitoid *T. chilonis* and its host *C. cephalonica* were obtained from the ICAR-National Bureau of Agricultural Insect Resources, Bengaluru, India, and Cryptox Biosolutions, Kanyakumari, India. The cultures were reared under laboratory conditions under ambient light and temperature. The rearing of both insects was based on the protocol published by the Directorate of Plant Protection, Quarantine & Storage, Department of Agriculture & Farmers Welfare, Ministry of Agriculture & Farmers Welfare, Government of India (Directorate of Plant Protection, Quarantine & Storage (Directorate of plant protection quarantine & storage & Government of India). *Corcyra cephalonica* was reared on a Sorghum and groundnut-based diet in plastic containers covered with muslin cloth. The moths emerging from these boxes were collected and transferred to breeding cages. The breeding cages were provided with 50% honey solution, and the eggs laid by the moths were collected and stored at 4°C. The parasitoids were reared on the UV-sterilised eggs of *C. cephalonica.* The sterilised eggs of *C. cephalonica* were pasted on paper cards (hereafter called egg cards) and placed in plastic boxes containing adult parasitoids for parasitisation. The adult parasitoids were also provided with a 50% honey solution.

## Experimental design

Egg cards for experiments were made by glueing 24-hour-old *C. cephalonica* eggs to a 1cm^2^ card and then sterilising the card to kill the host embryo. The size of egg cards for control and Resource Constant (RC) conditions, irrespective of parasitoid density, was fixed at 1cm^2^ (containing 500-600 eggs) to keep the resource density constant. However, for the Resource Abundant (RA) condition, the size of the card increases with increasing parasitoid density to avoid resource constraints, i.e. 2 x 1cm^2^, 4 x 1cm^2^ and 8 x 1cm^2^, respectively, for 2, 4 and 8 parasitoid densities (Fig. 1). A newly emerged female parasitoid mated within 8h was introduced into a 35x10mm Cell Culture Dish (Thermo Fisher Scientific, USA). 24 hours after their introduction, the experiment was conducted. All experiments were performed starting at 10:00 am (4 hours after light on) to minimise the temporal effect on parasitoid behaviour.

The egg cards were introduced into the Petri dish containing wasps for each experimental trial, and parasitisation was allowed for 2 hours, after which the wasps were removed. Since the 8 x 1 cm² egg cards for eight female densities in RA conditions are larger than the size of the 35 x 10 mm Cell Culture dishes, the wasps and egg cards for this treatment were placed in larger Petri dishes of size 90 x 14 mm (Tarsons Products Pvt. Ltd, India). After the experiment, the egg cards were stored at ambient room temperature during parasitoid development. The number of eggs parasitised in each Petri dish was counted four days after parasitisation, when the parasitised eggs turned from white to black. The wasp offspring emerged from the parasitised eggs within nine days of parasitisation. The number of offspring that emerged and their sex were recorded after 10 days.

## Resource utilisation analysis

The resource utilisation of parasitoids under competitive and non-competitive conditions was studied by comparing the number of eggs parasitised per female under different competitive conditions. To examine whether the presence of competitor(s) affects resource utilisation, the number of eggs parasitised per female at 2, 4 and 8 female densities was compared with the non-competitive one-female condition (n=78). We also compared the number of eggs parasitised per female between 2, 4 and 8 parasitoid densities to study the effect of parasitoid density on resource utilisation. Finally, we also analysed the impact of resource abundance on resource utilisation. The effect of resource abundance was studied under two resource conditions-resource-abundant (RA) and resource-constant (RC). The sample sizes used for the study are shown in Table 1.

**Table 1:**
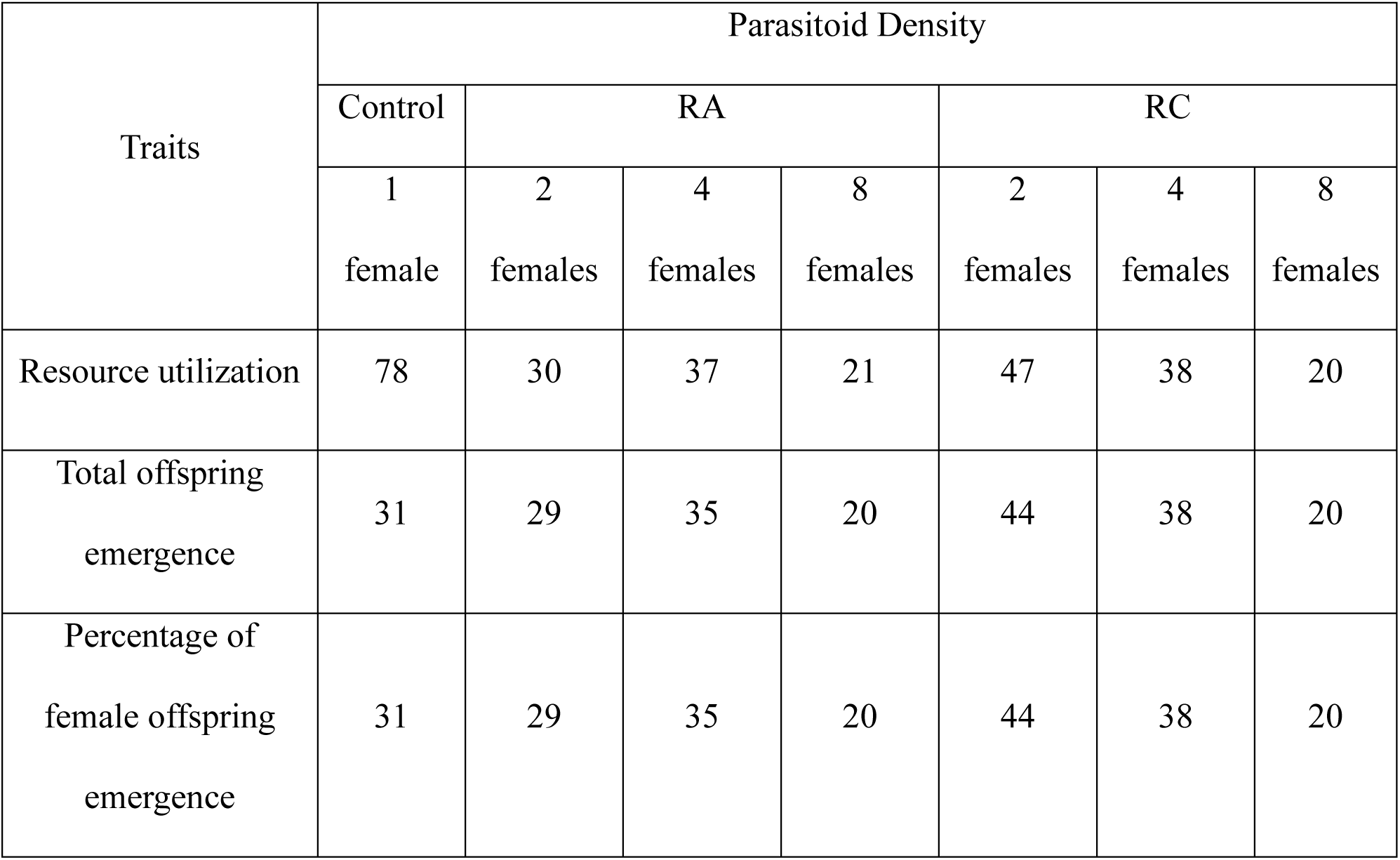
The number of replicate experimental trials for each measure of parasitoid performance at each parasitoid density and host density treatment. In the resource-abundant (**RA**) conditions, host egg availability increases with parasitoid density. In the resource-constrained condition (**RC**), the number of host eggs available does not increase with parasitoid density

## Percentage of total offspring and female emergence

The percentage of parasitised eggs from which the adult offspring emerged (percentage of total emergence) and the percentage of female offspring that emerged were used to represent juvenile mortality and sex ratio, respectively. The impact of the presence of a competitor on the emergence rate and offspring sex ratio was compared between competitive (2, 4, and 8 female density) and non-competitive one-female conditions (n = 31). The impact of parasitoid densities (2, 4 and 8 females) and the impact of two resource densities (RA and RC) on performance traits were also compared in this study. The sample sizes for both performance traits are given in Table 1.

## Statistical analysis

The number of eggs parasitised per female in the presence and absence of conspecifics was compared using the nonparametric Kruskal-Wallis Test and Dunn’s test. The effect of different resource abundances (RC vs RA) on resource utilisation (number of eggs parasitised) was compared using the Student’s t-test and Mann–Whitney U test. The effect of parasitoid density (2, 4 and 8 female densities) on the number of eggs parasitised per female was compared in the RA conditions using one-way ANOVA, and post hoc analysis was conducted using Tukey’s HSD Test. Meanwhile, the effect of parasitoid density (2, 4 and 8 female densities) on the number of eggs parasitised per female in the RC condition was compared using the Welch’s ANOVA.

The percentage of total emergence and female offspring that emerged from the parasitised eggs under different parasitoid densities was compared using the Kruskal-Wallis test. Dunn’s test was conducted as a post hoc analysis. Similarly, the effects of resource abundance on the emergence and sex ratio were compared using the Mann–Whitney U test and Welch’s t-test. The statistical analyses were done using R version 4.3.3 (R Core Team, 2024).

## Results

### Effect of competition on resource utilisation

The number of eggs parasitised per female was greater in the presence of conspecifics than in the absence of conspecifics under both resource density conditions (Fig. 2a, b; *p* <0.001 expect for 1 vs 8 females under RA conditions is *p* <0.05). In the RA condition, the number of eggs parasitised per female wasp at the 2 (18.5 ± 9.38), 4 (23.95 ± 8.98), and 8 (13.67 ± 6.84) female densities is significantly higher than in a one female condition (6.14 ± 10.18). Similarly, the number of eggs parasitised per female at 2 (20.79 ± 13.94), 4 (22.93 ± 7.17), and 8 (19.01 ± 6.61) female densities in the RC condition is also significantly higher than in the one female condition.

**Figure 2:**
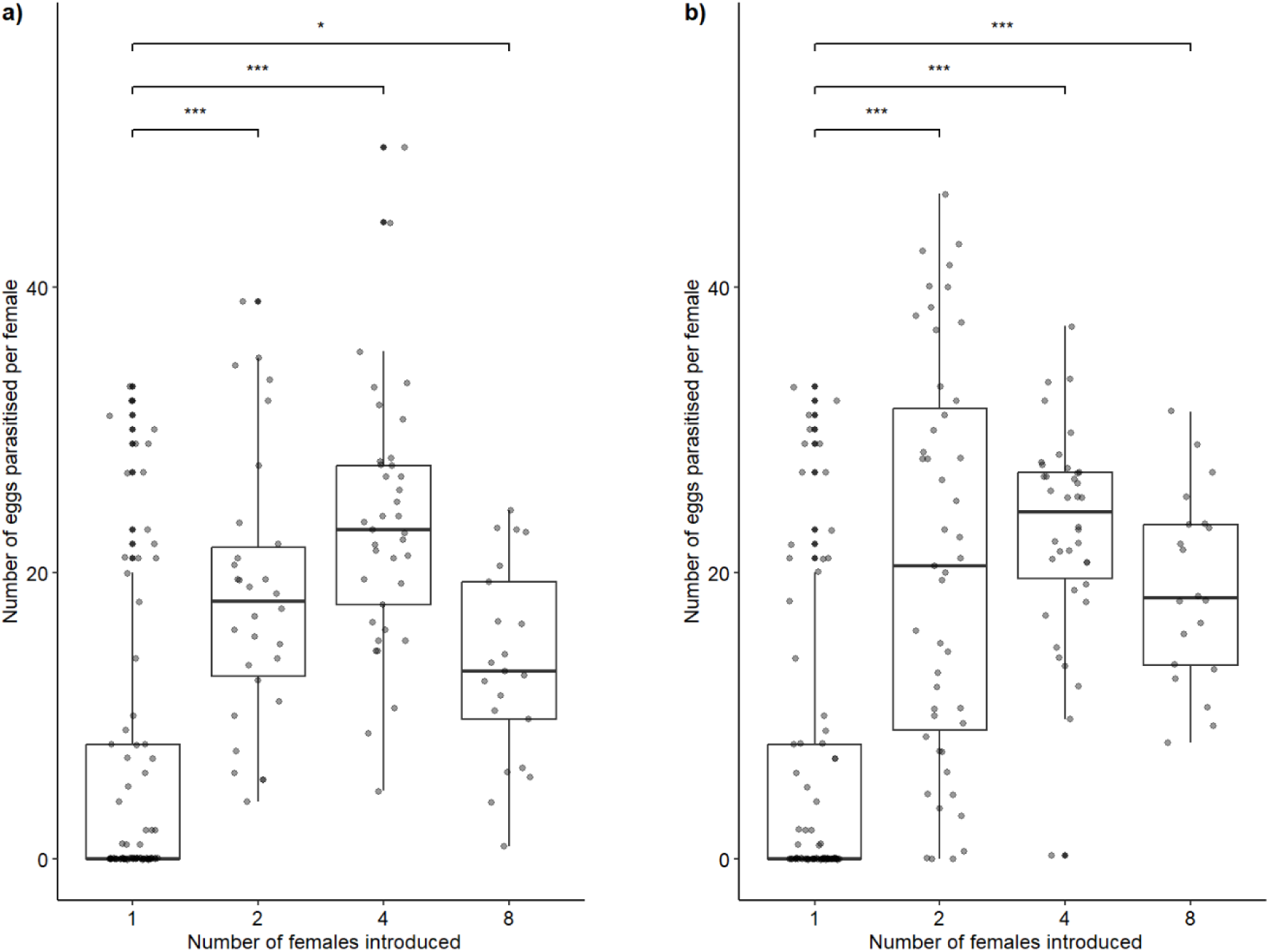
Resource utilisation in the presence and absence of conspecifics: The number of eggs parasitised per fem ale wasp in the non-competitive (1 female) condition and in the presence of increasing conspecifics in **RA (panel a)** and the **RC** condition **(Panel b)**. The number of asterisks indicates the level of significance. ‘*’ indicates marginal significance of ≤ 0.05 and ‘***’ indicates higher significance of ≤ 0.001.

Comparing the effect of parasitoid densities under competitive conditions (2, 4 and 8), in both RA and RC conditions, the number of eggs parasitised per female tended to be higher at the 4 female densities than in the two female and eight female densities (Fig. 3); however, this increase is significant only in the RA condition (*p-*value <0.0001) and not in the RC conditions (p-value = 0.12). We also found that resource density did not significantly affect resource utilisation in the overall comparison (*p* = 0.23). The overall mean resource utilisation in the RA condition is 19.47 ± 9.43; for RC, it is 21.23 ± 10.69. However, in pairwise comparison of the effect of resource density between different parasitoid densities, eight female densities under the RC condition have a significantly higher resource utilisation than the RA condition (*p* value for 2, 4 and 8, respectively, is 0.55, 0.74 and 0.01) (Fig. S1).

**Figure 3:**
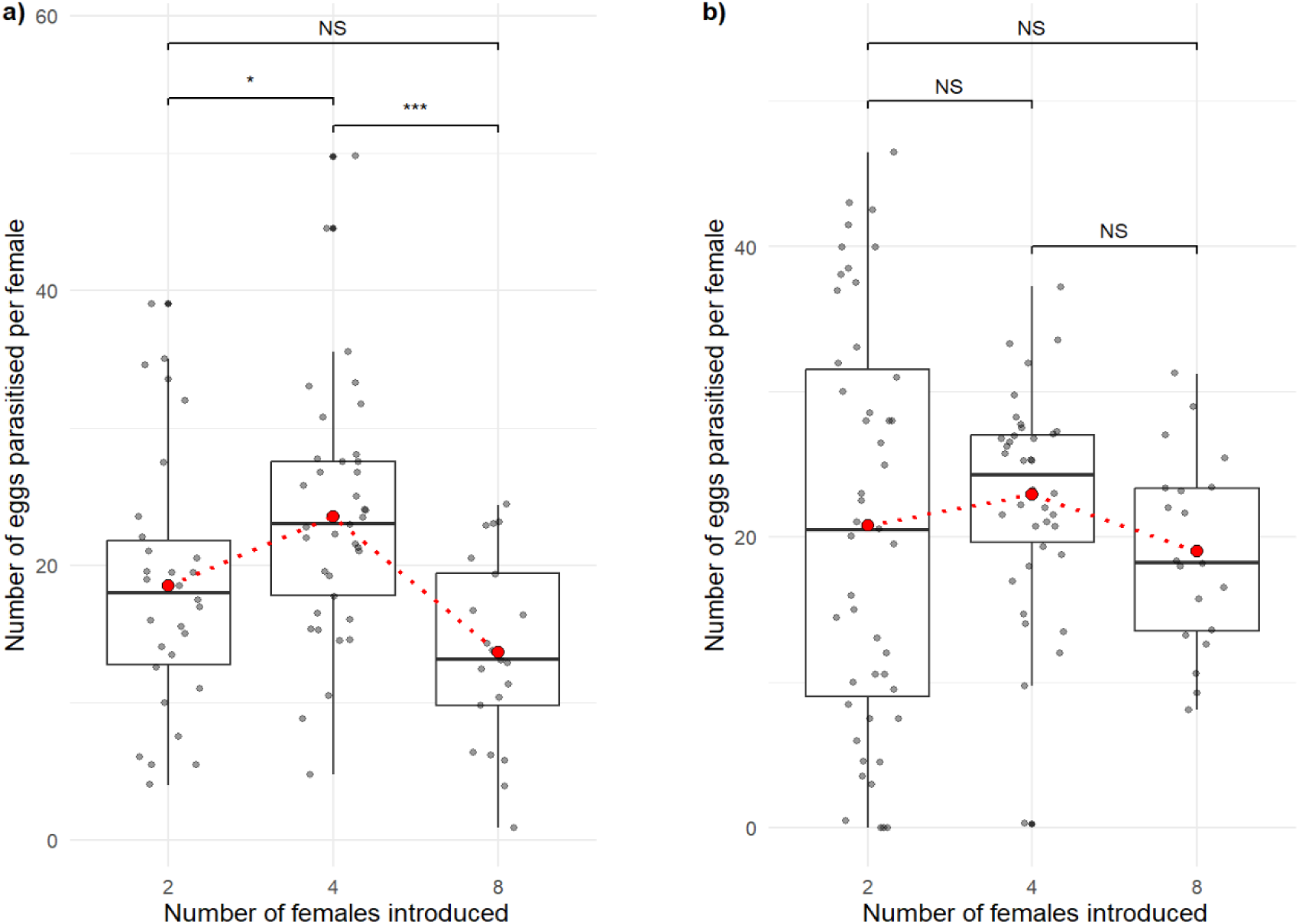
The number of eggs parasitised per female at varying parasitoid densities: The number of eggs parasitised per female at varying conspecific densities of 2, 4 and 8 in the **RA** condition (*p*-value <0.0001, **panel a**) and in the **RC** conditions (*p*-value = 0.12, **panel b**). After post-hoc analysis of RA *p*-value for 2-4 is 0.052, 2-8 is 0.12, and 4-8 <0.001. The number of asterisks indicates the level of significance; ‘*’ indicates marginal significance of ≤ 0.05, ‘***’ indicates higher significance of ≤ 0.001. NS indicates no significance. The red dotted trend line passes through the mean value.

### Effect of competition on the percentage of offspring emergence and secondary sex ratio

The percentage of offspring successfully emerging from parasitised eggs when there is only one female wasp was 75.5 ± 25.2. In the RA condition, the average percentage of offspring that emerged in two female, four female, and eight female conditions, respectively, is 73.2 ± 23.9, 60.5 ± 26.7, and 69.8 ± 20.7. For the RC condition, the average percentage of offspring that emerged under two females is 84.2 ± 16.9, under four females is 85 ±13.7, and under eight females is 87.5 ± 5.7 (Fig. 4).

**Figure 4:**
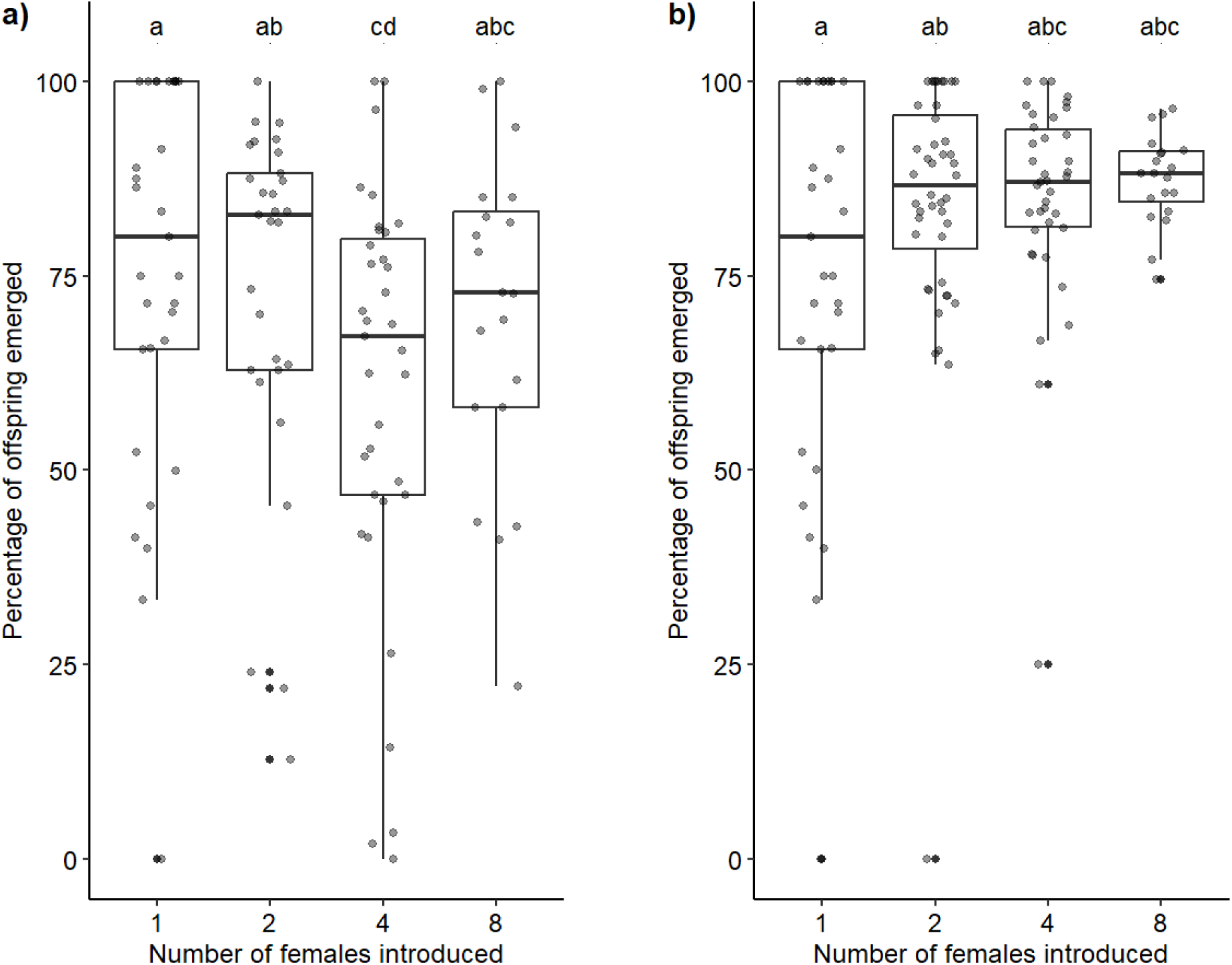
Percentage of offspring emerged under different parasitoid densities: The effect of the presence of conspecifics on the percentage of offspring that emerged in **RA (*p* value = 0.03; panel a)** and **RC (*p* value = 0.60; panel b)** conditions. The *p*-value of the percentage of offspring emergence under the RA condition after post hoc analysis for 1-2,1-4, 1-8, 2-4, 2-8, and 4-8, respectively, is 0.66, 0.04, 0.75, 0.14, 0.72, and 0.92. The similar superscript letters indicate no statistically significant difference between groups, while different letters show a significant difference at *p* ≤ 0.05.

Except at the density of four females, the offspring emergence rate under the RA condition was similar at all parasitoid densities (*p*-value = 0.03; Fig. 4a). Similarly, under RC conditions, emergence did not different between different parasitoid densities (*p*-value = 0.60; Fig. 4b). The overall percentage of offspring that emerged in the RA condition (67.1 ± 24.5) was lower than under the RC condition (85.1 ±14.1) (*p* <0.0001; Fig. 5a). This was true at each parasitoid densities (*p-*value for 2, 4 and 8, respectively 0.02, <0.0001 and 0.03; Fig. 5b).

**Figure 5:**
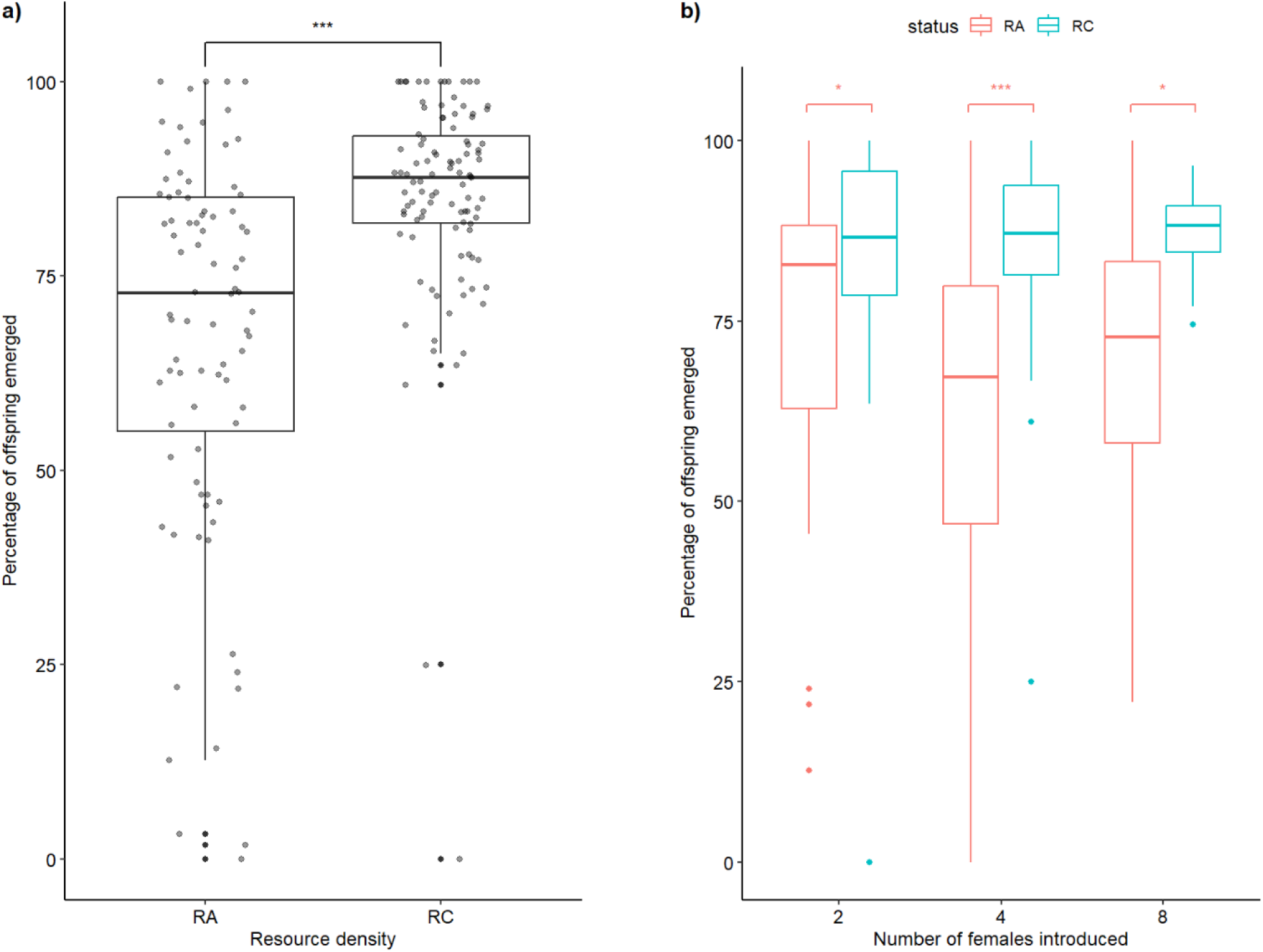
The effect of resource density on the percentage of offspring that emerged: The overall effect of resource density on percentage of offspring emerged **(panel a)** and the pairwise comparison across different parasitoid densities 2, 4 and 8 **(panel b)**. The number of asterisks indicates the level of significance; ‘*’ indicates significance of ≤0.05, ‘***’ indicates significance of ≤ 0.001.

There was an overall female-biased sex ratio among the offspring produced. On average, the percentage of female offspring was 68.75 ± 24.17 for an adult alone. This did not change with increasing number of conspecifics under either resource availability treatment (*p-*value for RA is 0.33 and for RC is 0.13; Fig. S2). Similarly, the secondary sex ratio did not differ between the RA and RC resource densities (*p-*value=0.57; Fig. S3).

### Mathematical model of progeny emergence rate under resource-constrained conditions

The pattern of increased emergence rate of adult progeny wasps under host resources-constrained conditions (RC) (Fig. 5) could be due to increased superparasitism at lower host egg density. Typically, only one *T. chilonis* individual emerges from a parasitised host (Chowdhury et al., 2016; Manisha et al., 2020). Under single-female conditions, the average emergence rate was approximately 75%, indicating that about 25% of parasitised eggs failed to yield adult progeny (Fig. 4). This suggests that parasitism was unsuccessful in roughly a quarter of cases. We hypothesise that superparasitism (multiple females laying eggs in the same host) of hosts that are among the failed 25% parasitism may increase the likelihood that at least one offspring successfully emerges. Consequently, this could raise the overall emergence rate. To investigate this possibility, we modelled the relationship between offspring emergence rate and the number of females foraging together, across a range of host egg densities. The model incorporated oviposition rates and survival (emergence) data from our observations.

Let there be *M* high-quality host eggs that are parasitized by *N* wasps with *p* being the probability of an individual host egg being parasitized by a single wasp. Then, the probability that a host is parasitized by *i* of the *N* parasitoids is given by the Binomial distribution:

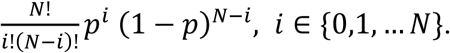

Plugging in 𝑖 = 0 gives probability that an egg escapes parasitism,

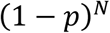

and yields the following average number of parasitized high-quality hosts (𝑖 ≥ 1)

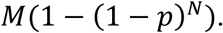

For a given number of adult female wasp 𝑁 ∈ {1,2,4,8}, the probability *p* is chosen to match the parasitized eggs per wasp seen in the data (Fig. 2), 7 parasitized eggs for a single wasp (𝑁 = 1) and 20 eggs per wasp for 𝑁 ∈ {2,4,8}.

Also based on the data (Fig. 4), we set the juvenile parasitoid survival probability inside the host egg to be 𝑠 = 0.75 when the egg is parasitized by a single wasp. Assuming a random model of juvenile wasp survival, an egg parasitized by *i* wasps will have adult offspring emergence with probability

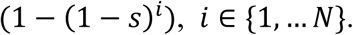

This gives the probability of adult wasp emergence from a parasitized egg to be 75% (as expected for 𝑖 = 1 when there are no superparasitism), and with superparasitism 94% (for 𝑖 = 2) and 98% (for 𝑖 = 3).

The fraction of parasitized eggs resulting in parasitoid emergence is now given by

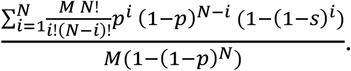

This ratio is plotted in Fig. 6 for 𝑁 ∈ {1,2,4,8} adult female foraging wasps, and a varying number of high-quality eggs. Recall from above that the probability *p* is dependent on *M* as the number of parasitized eggs per wasp is fixed. As shown in Fig. 6, increasing superparasitism driven by multiple wasps parasitizing a low number of high-quality eggs can enhance the percentage parasitoid emergence to up to 90% (𝑁 = 8*, M=200*) from its basal value of 75% (𝑁 = 1) which is similar to what we found in the empirical study with resource constraint (Figs. 5 and 6).

**Figure 6:**
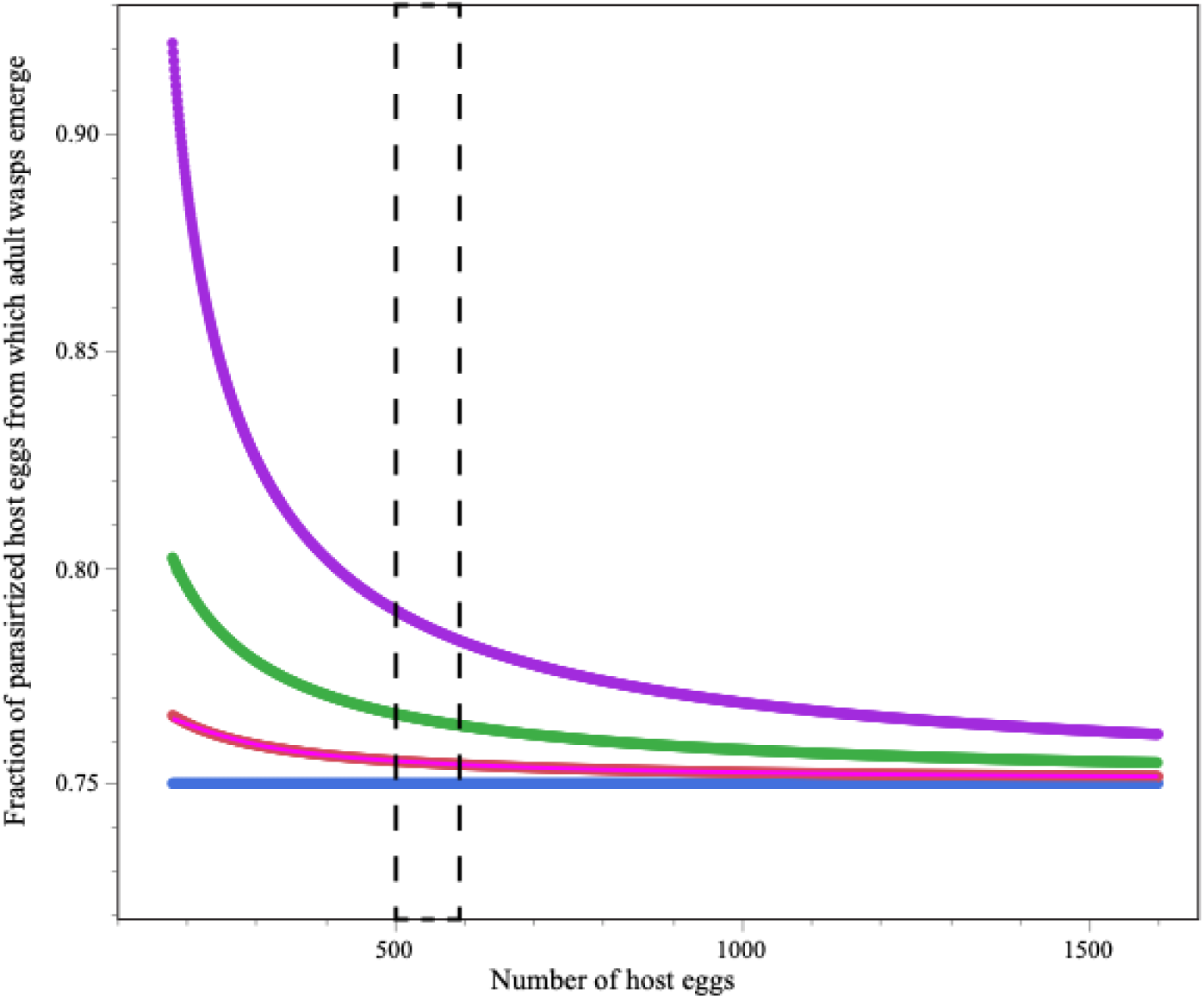
The association of fraction of parasitized eggs yielding adult parasitoids with host egg availability in the model. The coloured curves represent different numbers of foraging wasps (1-blue, 2-pink, 4-green, 8-purple). The dashed rectangle marks the resource constrained conditions in the experimental study.

## Discussion

Contrary to theoretical predictions (Robert et al., 2016), we found that the egg parasitoid *T. chilonis*, under intraspecific exploitative competition, parasitised more eggs in the presence of conspecifics than when alone. With increasing parasitoid densities, the individual rate of parasitism shows a similar trend of increasing at four female densities and a decline at eight female densities at both resource conditions. However, the resource density did not affect the number of eggs parasitised per female. The fraction of eggs parasitised from which progeny successfully emerged increased at low resource density. We made a simple mathematical model to demonstrate that at low resource density, superparasitism can lead to high population level emergence. The secondary sex ratio (percentage of female offspring emerging) was neither affected by the presence of conspecifics nor by the density of parasitoids, but showed a marginal trend of declining with increasing parasitoid density in resource constant conditions.

Previous studies of other parasitoid species have shown that the foraging decisions are affected by the presence of conspecific competitors (Ives, 1989; Visser et al., 1999; Harvey et al., 2013). Yet, the direction of this effect differs based on the competing species. For instance, studies on clutch size decisions of the solitary parasitoid *Comperiella bifasciata* Howard (Hymenoptera: Encyrtidae) show that females that were kept together with conspecifics before parasitisation laid more eggs than females kept alone (Visser & Rosenheim, 1998) However females of the gregarious parasitoid *Aphaereta minuta* Nees (Hymenoptera: Braconidae), kept with conspecifics before parasitisation produced smaller clutches than those kept alone (Visser, 1996). The patch allocation time, which is a component of the foraging decision of parasitoids, is also influenced by the presence of conspecifics and has been found to increase or decrease with competition (Visser et al., 1992). The parasitoid *Hyposoter horticola* Gravenhorst (Hymenoptera: Ichneumonidae) re-visits patches more frequently in the presence of a conspecific competitor than in the absence of a competitor (Couchoux & van Nouhuys, 2014). On the other hand, the pupal parasitoid *P. vindemmiae* Rondani (Hymenoptera: Pteromalidae) spent less time in patches when the number of competitors increased (Goubault et al., 2005). The effect of the presence of conspecifics on resource utilisation also differs among non-aggressive parasitoids related to those used in our study. In the case of *Trichogramma pintoi* Voegelé and *T. minutum* Riley (Hymenoptera: Trichogrammatidae), *T. pintoi* shortened patch residence time under competition, whereas *T. minutum* did not modify patch residence time based on competition, leading Robert et al (2016) to conclude that resource utilisation of adult parasitoids towards the presence of a competitor is parasitoid species-specific. Thus, we also conclude that the response of *T. chilonis* to resource utilisation in the presence of a conspecific might be a species-specific response.

One possible reason for increased resource utilisation is to compensate for the potential decline in individual fitness due to superparasitism. The ability of *T. chilonis* to distinguish between parasitised and unparasitized eggs varies with their previous ovipositional experiences. Those with no ovipositional experience tend to parasitise unparasitized eggs or previously parasitised eggs with the same probability (Miura et al., 1994). This suggests that the observed increase in resource utilisation may be an adaptive strategy, allowing *T. chilonis* to maximise individual reproductive success despite potential loss due to superparasitism.

While the presence of a conspecific significantly increased the number of eggs parasitised per female, increasing parasitoid density beyond two did not further increase their resource utilisation. There was an upward trend in parasitism as female density increased from two to four, followed by a decline at eight at both resource densities. Contrary to what we found, Yan *et al*. (2023) found a decreasing trend in parasitism per female *T. pintoi,* with increasing parasitoid densities (Yan et al., 2023). Similarly, Kfir, (1981) found that per female parasitism by *T. pretiosum* declined when parasitoid density increased from 2 to 4, but did not significantly vary when the parasitoid density increased from 4 to 16 (Kfir, 1981). In *T. chilonis,* the decline in the number of eggs parasitised per female is highly significant only between four and eight female densities in the RA condition. One of the reasons for the very low parasitisation under 8-female density in the RA condition could be due to the increase in the spatial scale of the enclosure made to accommodate a large number of egg cards under resource abundance. For *Trichogramma*, differences in the temporal and spatial scale distribution of resources can affect their host finding success (Burte et al., 2023). Overall, we conclude that increasing parasitoid density beyond two females does not significantly affect resource utilisation by *T. chilonis*. While it remains uncertain whether female wasps perceive conspecific presence through visual, olfactory, or tactile senses (Pexton & Mayhew, 2005), our study suggests that *T. chilonis* may not detect the number of conspecifics around them, at least up to eight females, but can recognise when at least one conspecific is in the vicinity, likely through odour cues (Gonthier et al., 2023).

The number of eggs parasitised per female was the same when offered abundant (RA) or constant (RC) resource density. A previous study of the functional response of *T. chilonis* on *C. cephalonica* found that *T. chilonis* follows Hollings type II functional response (Qin et al., 2023). In parasitoids following type II functional response, the individual parasitisation rate increases at lower host densities, while it decreases at higher host densities (Holling, 1966; Tazerouni et al., 2019). For another *Trichogramma* parasitoid, which follows type II functional response, *T. pretiosum,* above the host density of 300 eggs, the number of eggs parasitised per female reaches a plateau in the functional response curve in all 2, 4 and 8 parasitoid densities (Kfir, 1983). It is possible that we did not find an effect of resource density because the density of eggs was high (500-600 eggs per 1 cm^2^ card) even under the resource-constrained condition. However, since there was an increase in resource use in response to the presence of conspecifics, it may be that the wasps use the presence of competitors, rather than the density of host eggs, as a signal of declining host availability (van Alphen & Visser, 1990).

One outcome of competition for parasitoids is superparasitism, in which a mother lays eggs in a previously parasitised host. This can lead to the mortality of offspring if only one can develop. We found that neither the presence of a competitor nor the higher parasitoid density affects the survival of offspring (percentage emergence). However, emergence from parasitized eggs was higher when resources were constrained than when resources were abundant. As resource density decreases, we expect the rate of superparasitism to increase (Kraft & Van Nouhuys, 2013). While generally superparasitism is detrimental at the individual level for solitary parasitoids, it can be adaptive at the population level since it will increase the probability of at least one offspring emerging from a parasitised host (Mackauer & Chau, 2001). This may be true in our system, where not all parasitism leads to successful parasitoid development even when a host is singly parasitised, and the wasps are foraging simultaneously. Using a simple probabilistic model, we demonstrated that if the foraging parasitoids do not avoid superparasitism, the emergence rate is high when multiple wasps forage. It also decreases with increasing host density as superparasitism decreases. The resource-constrained condition with multiple foraging females qualitatively matches the model. This suggests that the wasps may not be avoiding superparasitism perhaps because the wasps are still naïve. While individual success declines with superparasitism, at the population level it results in a high rate of parasitism, which would be advantageous for biological control.

Parasitoid wasps are Hymenoptera, and thus have a haplodiploid sex determination system which allows mated females to control the sex of their offspring. Many exhibit a female-biased sex ratio because, where resources are sufficient, it is advantageous to have more daughters (Somjee et al., 2011). In our study, there was a female bias, which was not affected by the given resource densities, suggesting no perceived variation in host quality. There was, however, a marginal trend of declining percentage of female offspring at higher parasitoid densities in resource constant conditions. According to Local Mate Competition theory, the proportion of male offspring produced should increase with the number of foundress females visiting a patch, and the offspring sex ratio should change from female-biased toward a sex ratio of 1:1 (Hamilton, 1967). For instance, *Trichogramma pretiosum* (Luck et al., 2001) and *T. evanescens* (Waage & Lane, 1984) show lower female sex bias with increasing foundress number. Two other *Trichogramma* species, *T. minutum* and *T. pintoi*, produced more male offspring when the parasitoid density was ten females in a patch than when they are alone (Martel & Boivin, 2004). In *T. pintoi*, the percentage of male offspring increased at a maternal density greater than four (Yan *et al*., 2023). In our study, the marginal trend that we observed at higher founder density in a resource constant condition indicates that *T. chilonis* might also follow the male-biased sex ratio under more competitive conditions.

Understanding the effect of competition on resource utilisation and other performance traits gives us insights into the population dynamics of parasitoids, as well as pest control efficiency (Courchamp et al., 1999; Bográn et al., 2002; Ode & Hardy, 2008; Bompard et al., 2013). Taken along with previously studied parasitoids exhibiting exploitative competition, it is evident that the presence of competitors can have both positive and negative effects on resource utilisation and other performance traits, which differ between parasitoid species. One of the reasons is that it is difficult to come up with a pattern is because there are few documented studies of exploitative competition among parasitoids. In our study, we found that in the presence of conspecifics, *T. chilonis* increases its resource utilisation. Similarly, the percentage of offspring that survive to emerge is higher when resources are more limited. Thus, *T. chilonis* can perform relatively well under intraspecific competitive conditions. This might be one of the reasons for the high performance of *T. chilonis* in augmentative biological control against many pest species (Sharma et al., 2020; Li et al., 2024)

## Acknowledgment

We acknowledge Prof. Kavita Iswaran for her suggestions on statistical analysis, Tapasya Tapa and Shreya Gangwal for their assistance in rearing the insect culture, and the Population and Community Ecology lab members for their ideas and discussions. PM acknowledges the National Postdoctoral Fellowship from the Anusandhan National Research Foundation for funding the research, as well as the Centre for Ecological Sciences, Indian Institute of Science, for providing the facility. PM also thanks Prajit J, Central University of Kerala, for the suggestions and discussions.

## Supplementary figures for the paper

**Fig. S1:**
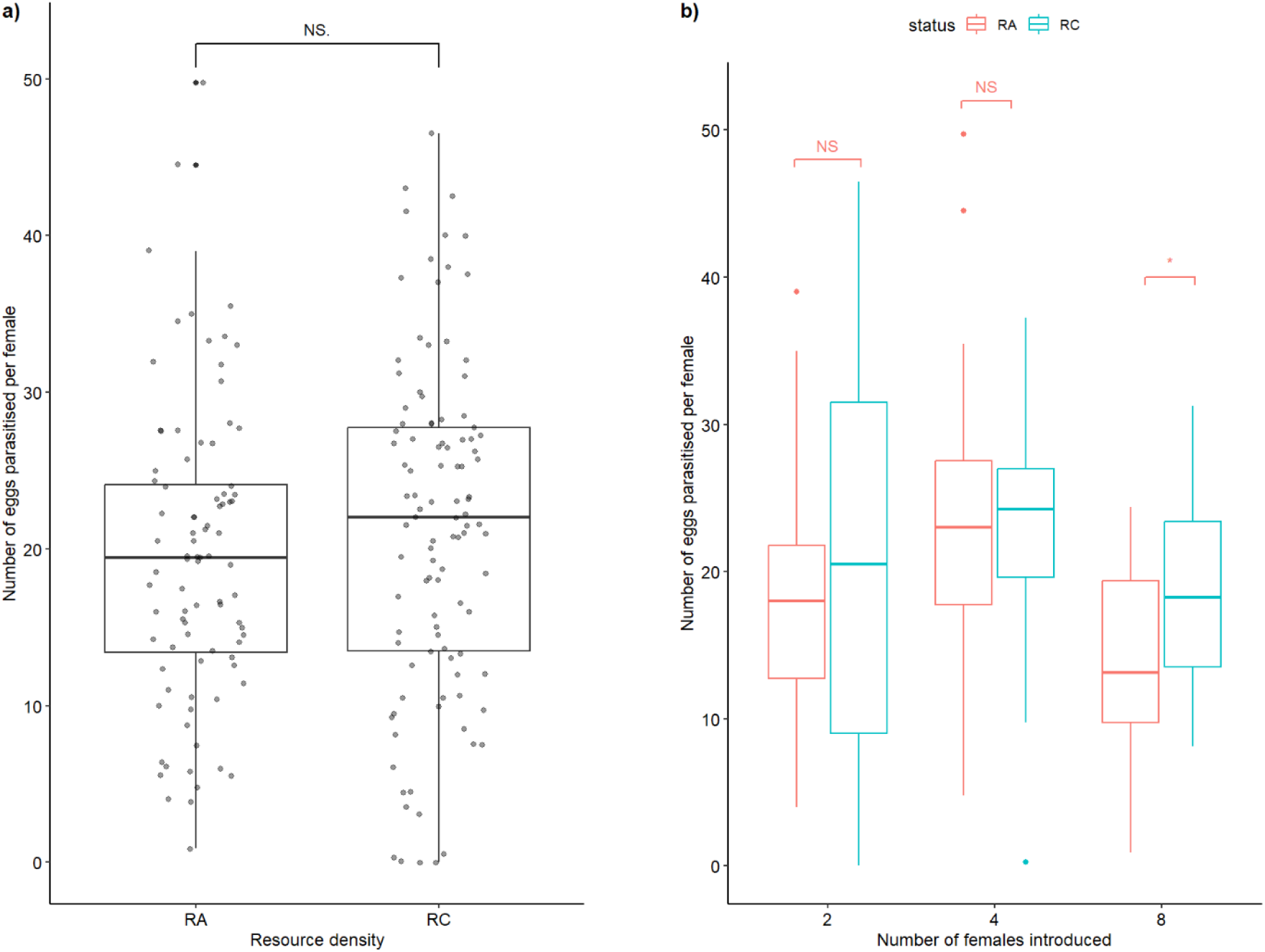
The comparison between the number of eggs parasitised by a female in two resource densities: Overall comparison between RA and RC conditions **(panel a)** and pairwise comparison across parasitoid densities of 2 (*p* value= 0.55), 4 (*p* value= 0.74), and 8 *(p* value = 0.01) under RA (red) and RC (blue) conditions (**panel b**). The number of asterisks indicates the level of significance; ‘*’ indicates significance of ≤ 0.05, NS indicates no significance.

**Fig. S2:**
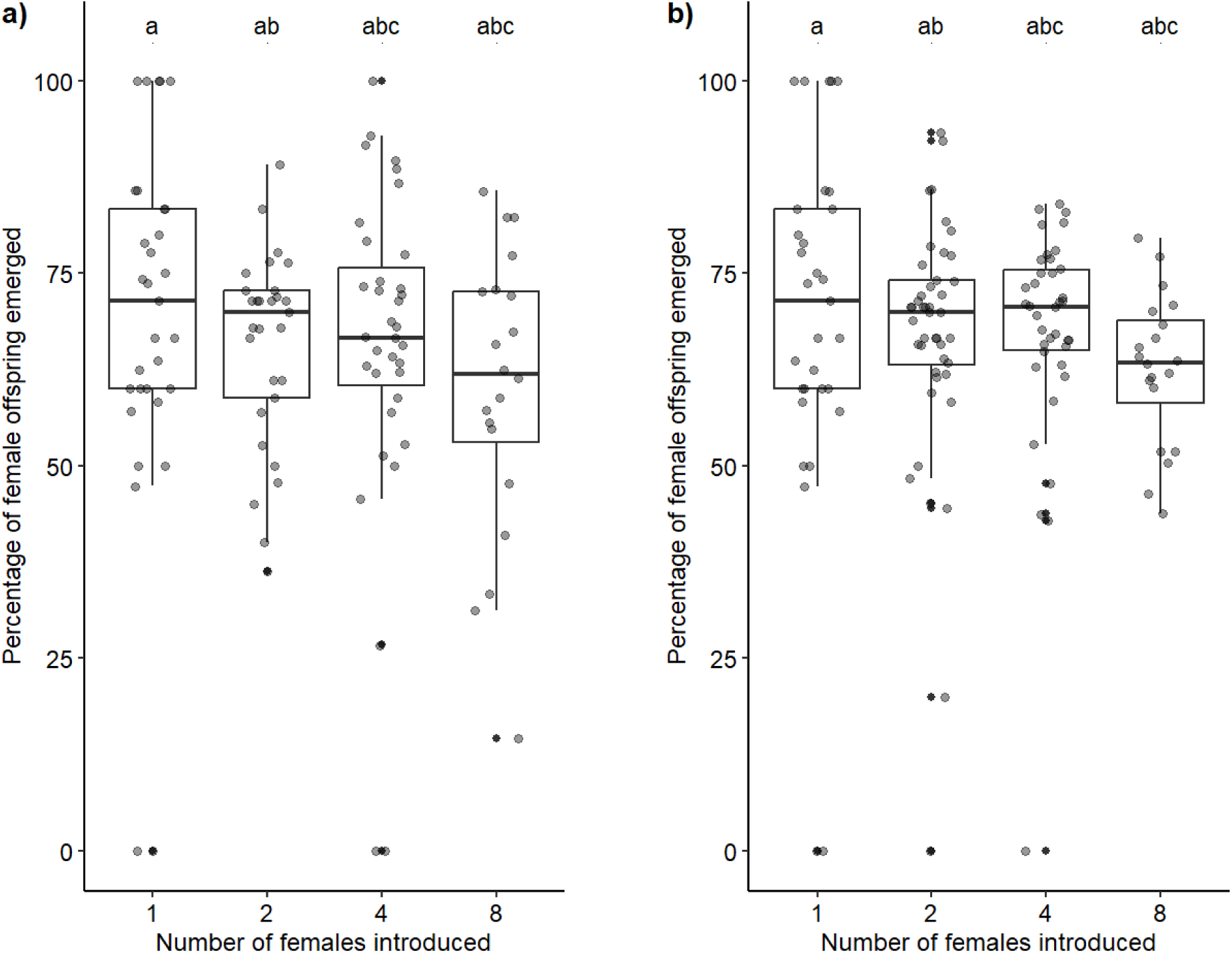
The offspring sex ratio at various parasitoid densities: The offspring sex ratio under **RA (panel a)** and **RC (panel b)** conditions. The p-value for RA is 0.33 and for RC is 0.13. The similar superscript letters indicate no statistically significant difference between groups.

**Fig. S3:**
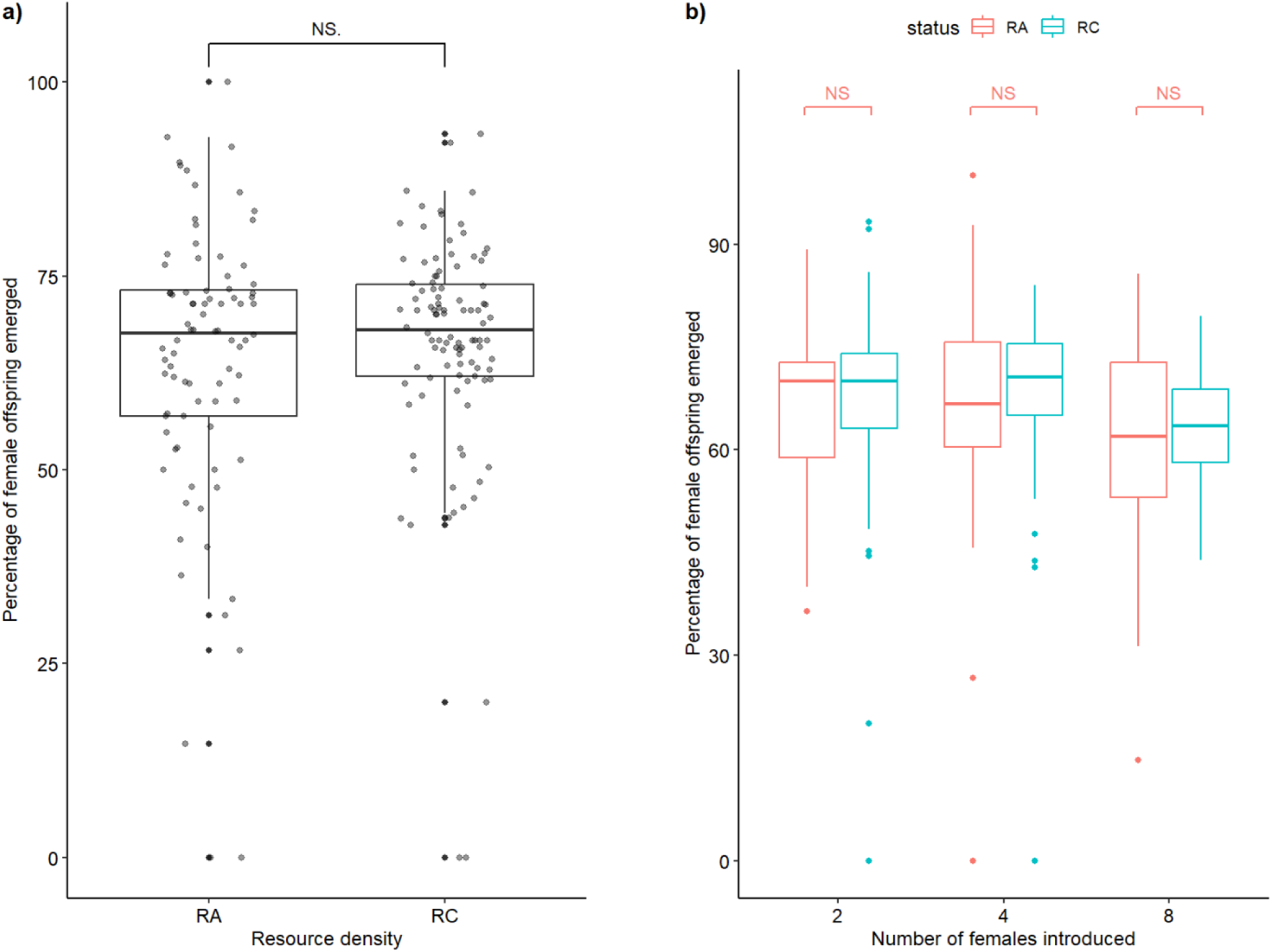
The effect of resource density on offspring sex ratio: Overall comparison between RA and RC conditions **(panel a)** and pairwise comparison across parasitoid densities **(panel b)** on the effect of resource densities on the offspring secondary sex ratio.

## References

Agnew P, Hide M, Sidobre C & Michalakis Y (2002) A minimalist approach to the effects of density-dependent competition on insect life-history traits. Ecological Entomology 27:396–402.

van Alphen JJM & Visser ME (1990) Superparasitism as an Adaptive Strategy for Insect Parasitoids. Annual Review of Entomology 35:59–79.

Batchelor TP, Hardy ICW, Barrera JF & Pérez-Lachaud G (2005) Insect gladiators II: Competitive interactions within and between bethylid parasitoid species of the coffee berry borer, *Hypothenemus hampei* (Coleoptera: Scolytidae). Biological Control 33:194–202.

Bernhardt JR, Kratina P, Pereira AL, Tamminen M, Thomas MK & Narwani A (2020) The evolution of competitive ability for essential resources. Philosophical Transactions of the Royal Society B: Biological Sciences 375:20190247.

Bográn CE, Heinz KM & Ciomperlik MA (2002) Interspecific competition among insect parasitoids: Field experiments with whiteflies as hosts in cotton. Ecology 83:653–668.

Bompard A, Amat I, Fauvergue X & Spataro T (2013) Host-parasitoid dynamics and the success of biological control when parasitoids are prone to Allee effects. PLoS ONE 8:e76768.

Brodeur J & Boivin G (2004) Functional ecology of immature parasitoids. Annual Review of Entomology 49:27–49.

Burte V, Cointe M, Perez G, Mailleret L & Calcagno V (2023) When complex movement yields simple dispersal: behavioural heterogeneity, spatial spread and parasitism in groups of micro-wasps. Movement Ecology 11:13.

Chowdhury Z, Alam S, Dash C, Maleque M & Akhter A (2016) Determination of parasitism efficacy and development of effective field release technique for *Trichogramma* spp. (Trichogrammatidae: Hymenoptera). American Journal of Experimental Agriculture 10:1–7.

Cody M, MacArthur R & Diamond J (1975) Ecology and evolution of communities. Harvard University Press.

Couchoux C & van Nouhuys S (2014) Effects of intraspecific competition and host-parasitoid developmental timing on foraging behaviour of a parasitoid wasp. Journal of Insect Behavior 27:283–301.

Courchamp F, Clutton-Brock T & Grenfell B (1999) Inverse density dependence and the Allee effect. Trends in Ecology & Evolution 14:405–410.

Cusumano A, Peri E, Boivin G & Colazza S (2015) Fitness costs of intrinsic competition in two egg parasitoids of a true bug. Journal of Insect Physiology 81:52–59.

Cusumano A, Peri E & Colazza S (2016) Interspecific competition/facilitation among insect parasitoids. Current Opinion in Insect Science 14:12–16.

Directorate of plant protection quarantine & storage & Government of India Mass production protocol of bio-agents.

Forsman JT, Seppänen J-T & Mönkkönen M (2002) Positive fitness consequences of interspecific interaction with a potential competitor. Proceedings of the Royal Society of London. Series B: Biological Sciences 269:1619–1623.

Gonthier J, Zhang YB, Zhang GF, Romeis J & Collatz J (2023) Odor learning improves efficacy of egg parasitoids as biocontrol agents against Tuta absoluta. Journal of Pest Science 96:105–117.

Goubault M, Outreman Y, Poinsot D & Cortesero AM (2005) Patch exploitation strategies of parasitic wasps under intraspecific competition. Behavioral Ecology 16:693–701.

Hamilton WD (1967) Extraordinary Sex Ratios. Science 156:477–488.

Harvey JA, Gols R & Strand MR (2009) Intrinsic competition and its effects on the survival and development of three species of endoparasitoid wasps. Entomologia Experimentalis et Applicata 130:238–248.

Harvey JA, de Haan L, Verdeny-Vilalta O, Visser B & Gols R (2019) Reproduction and offspring sex ratios differ markedly among closely related hyperparasitoids living in the same microhabitats. Journal of Insect Behavior 32:243–251.

Harvey JA, Poelman EH & Tanaka T (2013) Intrinsic inter-and intraspecific competition in parasitoid wasps. Annual Review of Entomology 58:333–351.

Hasegawa K & Yamamoto S (2009) Effects of competitor density and physical habitat structure on the competitive intensity of territorial white spotted charr *Salvelinus leucomaenis*. Journal of Fish Biology 74:213–219.

Holling CS (1966) The functional response of invertebrate predators to prey density. Memoirs of the Entomological Society of Canada 98:5–86.

Ives AR (1989) The optimal clutch size of insects when many females oviposit per patch. Source: The American Naturalist 133:671–687.

Johnson CA, Grant JWA & Giraldeau LA (2004) The effect of patch size and competitor number on aggression among foraging house sparrows. Behavioral Ecology 15:412–418.

Kaplan I & Denno RF (2007) Interspecific interactions in phytophagous insects revisited: A quantitative assessment of competition theory. Ecology Letters 10:977–994.

Kfir R (1981) Effect of hosts and parasite density on the egg parasite *Trichogramma pretiosum* [Hym.: Trichogrammatidae]. Entomophaga 26:445–451.

Kfir R (1983) Functional response to host density by the egg parasite *Trichogramma pretiosum*. Entomophaga 28:345–353.

King BH (1993) Sex-ratio manipulation by parasitoid wasps. Evolution and diversity of sex ratio in insects and mites. (ed by Wrensch DL & M Ebbert) Chapman and Hall, New York, London, pp 418–441.

King BH & Seidl SE (1993) Sex ratio response of the parasitoid wasp *Muscidifurax raptor* to other females. Oecologia 94:428–433.

Kishani Farahani H, Moghadassi Y, Alford L & van Baaren J (2019) Effect of interference and exploitative competition on associative learning by a parasitoid wasp: a mechanism for ideal free distribution? Animal Behaviour 151:157–163.

Koutsidi M, Lazaris A, Peristeraki P, Tserpes G & Tzanatos E (2024) Quantification of intraspecific and interspecific competition in fish species of the Aegean Sea. ICES Journal of Marine Science 81:334–347.

Kraft TS & Van Nouhuys S (2013) The effect of multi-species host density on superparasitism and sex ratio in a gregarious parasitoid. Ecological Entomology 38:138–146.

Li X, Chen T, Chen L, Ren J, Ullah F, Yi S, Pan Y, Zhou S, Guo W, Fu K, Li YX & Lu Y (2024) *Trichogramma chilonis* is a promising biocontrol agent against *Tuta absoluta* in China: Results from laboratory and greenhouse experiments. Entomologia Generalis 44:357–365.

Luck RF, Janssen JAM, Pinto JD & Oatman ER (2001) Precise sex allocation, local mate competition, and sex ratio shifts in the parasitoid wasp *Trichogramma pretiosum*. Behavioral Ecology and Sociobiology 49:311–321.

Mackauer M & Chau A (2001) Adaptive self superparasitism in a solitary parasitoid wasp: The influence of clutch size on offspring size. Functional Ecology 15:335–343.

Manisha BL, Visalakshi MM, Sairam Kumar D V. & Kishore Varma P (2020) Resource efficient and cost-reduction technology for *Trichogramma chilonis* ishii (Hymenoptera: Trichogrammatidae) production. Journal of Biological Control 34:43–46.

Martel V & Boivin G (2004) Impact of competition on sex allocation by *Trichogramma*. Entomologia Experimentalis et Applicata 111:29–35.

Miura K & Kobayashi M (1998) Effects of host-egg age on the parasitism by *Trichogramma chilonis* Ishii (Hymenoptera : Trichogrammatidae), an egg parasitoid of the diamondback moth. Applied Entomology and Zoology 33:219–222.

Miura K, Matsuda S & Kobayashi M (1994) Discrimination between parasitized and unparasitized hosts in an egg parasitoid, *Trichogramma chilonis* Ishii (Hymenoptera: Trichogrammatidae). Applied Entomology and Zoology 29:317–322.

Nakamatsu Y, Harvey JA & Tanaka T (2009) Intraspecific competition between adult females of the hyperparasitoid *Trichomalopsis apanteloctena* (Hymenoptera: Chelonidae), for Domination of *Cotesia kariyai* (Hymenoptera: Braconidae) cocoons. Ann. Entomol. Soc. Am 102:172–180.

Ode PJ & Hardy ICW (2008) Parasitoid sex ratios and biological control. Behavioral Ecology of Insect Parasitoids. (ed by É Wajnberg, C Bernstein & J van Alphen) Wiley, pp 253–291.

Ode PJ, Vyas DK & Harvey JA (2022) Extrinsic inter- and intraspecific competition in parasitoid wasps. Annual Review of Entomology 67:305–328.

Pekkonen M, Ketola T & Laakso JT (2013) Resource availability and competition shape the evolution of survival and growth ability in a bacterial community. PLoS ONE 8:e76471.

Pexton JJ & Mayhew PJ (2005) Clutch size adjustment, information use and the evolution of gregarious development in parasitoid wasps. Behavioral Ecology and Sociobiology 58:99–110.

Qin Z, Li D, Luo Y & Cuthbertson AGS (2023) Impact of different *Trichogramma chilonis* populations from sugarcane fields on parasitism of *Corcyra cephalonica*. Journal of Applied Entomology 147:819–824.

R Core Team (2024) R: A language and environment for statistical computing, R foundation for statistical computing, Vienna, Austria. https://www.R-project.org.

Richter-Boix A, Llorente GA & Montori A (2004) Responses to competition effects of two anuran tadpoles according to life-history traits. Oikos 106:39–50.

De Rijk M, Zhang X, Van Der Loo JAH, Engel B, Dicke M & Poelman EH (2016) Density-mediated indirect interactions alter host foraging behaviour of parasitoids without altering foraging efficiency. Ecological Entomology 41:562–571.

Riley LA & Dybdahl MF (2015) The roles of resource availability and competition in mediating growth rates of invasive and native freshwater snails. Freshwater Biology 60:1308–1315.

Robert FA, Brodeur J & Boivin G (2016) Patch exploitation by non-aggressive parasitoids under intra- and interspecific competition. Entomologia Experimentalis et Applicata 159:92–101.

Sharma S, Shera PS, Kaur R & Sangha KS (2020) Evaluation of augmentative biological control strategy against major borer insect pests of sugarcane—a large-scale field appraisal. Egyptian Journal of Biological Pest Control 30:127.

Singhamuni SAA, Jayasuriya MIUF, Hemachandra KS & Sirisena UGAI (2015) Evaluation of the potential of *Trichogramma chilonis* Ishii (Hymenoptera: Trichogrammatidae) as a Bio-control Agent for *Trichoplusia ni*, Cabbage Semi-looper. Tropical Agricultural Research 26:223–236.

Somjee U, Ablard K, Crespi B, Schaefer PW & Gries G (2011) Local mate competition in the solitary parasitoid wasp *Ooencyrtus kuvanae*. Behavioral Ecology and Sociobiology 65:1071–1077.

Tao Z, Shen C, Qin W, Nie B, Chen P, Wan J, Zhang K, Huang W & Siemann E (2024) Fluctuations in resource availability shape the competitive balance among non-native plant species. Ecological Applications 34:e2795.

Tazerouni Z, Talebi AA & Rezaei M (2019) Functional response of parasitoids: Its impact on biological control. Parasitoids: biology, behavior and ecology. (ed by E Donnelly) Nova Science Publishers, New York, pp 1–119.

Than AT, Ponton F & Morimoto J (2020) Integrative developmental ecology: a review of density-dependent effects on life-history traits and host-microbe interactions in non-social holometabolous insects. Evolutionary Ecology 34:659–680.

Visser ME (1996) The influence of competition between foragers on clutch size decisions in an insect parasitoid with scramble larval competition. Behavioral Ecology 7:109–114.

Visser ME, van Alphen JJM & Nell HW (1992) Adaptive superparasitism and patch time allocation in solitary parasitoids : the influence of pre-patch experience. Behavioral Ecology and Sociobiology 31:163–171.

Visser ME, Jones TH & Driessen G (1999) Interference among insect parasitoids: a multi-patch experiment. Journal of Animal Ecology 68:108–120.

Visser B, Le Lann C, Snaas H, Hardy ICW & Harvey JA (2014) Consequences of resource competition for sex allocation and discriminative behaviors in a hyperparasitoid wasp. Behavioral Ecology and Sociobiology 68:105–113.

Visser ME & Rosenheim JA (1998) The Influence of competition between foragers on clutch size decisions in insect parasitoids. Biological Control 11:169–174.

Waage JK & Lane JA (1984) The reproductive strategy of a parasitic wasp: II. Sex allocation and local mate competition in *Trichogramma evanescens*. Journal of Animal Ecology 53:417–426.

Wajnberg E, Fauvergue X & Pons O (2000) Patch leaving decision rules and the Marginal Value Theorem: An experimental analysis and a simulation model. Behavioral Ecology 11:577–586.

Wang S & Callaway RM (2021) Plasticity in response to plant–plant interactions and water availability. Ecology 102:e03361.

Wang D, Lü L, He Y, Shi Q, Tu C & Gu J (2016) Mate choice and host discrimination behavior of the parasitoid *Trichogramma chilonis*. Bulletin of Entomological Research 106:530–537.

Wauters LA, Mazzamuto MV, Santicchia F, Van Dongen S, Preatoni DG & Martinoli A (2019) Interspecific competition affects the expression of personality-traits in natural populations. Scientific Reports 9:11189.

Xu HY, Yang NW & Wan FH (2013) Competitive interactions between parasitoids provide new insight into host suppression. PLoS ONE 8:e82003.

Yan Z, Yue JJ & Zhang YY (2023) Biotic and abiotic factors that affect parasitism in *Trichogramma pintoi* (Hymenoptera: Trichogrammatidae) as a biocontrol agent against *Heortia vitessoides* (Lepidoptera: Pyralidae). Environmental Entomology 52:301–308.

